# Ten species comprise half of the bacteriology literature, leaving most species unstudied

**DOI:** 10.1101/2025.01.04.631297

**Authors:** Paul A. Jensen

## Abstract

Microbiology research has historically focused on a few species of model organisms. Our bibliographic analysis finds extreme bias in the distribution of bacteriology research across species, with half of all papers referencing only ten species and 74% of all known species remaining unstudied. Microbiologists will need to broaden their perspective and embrace complexity to develop a complete understanding of the microbial world.

---

The microbiome revolution moved the goalposts for microbiologists. For centuries, microbiology has focused intensely on understanding a relatively small number of microbes. These model species were selected for their importance to health, the environment, industry, or simply because the species were easy to work with. Microbiologists maintained their focus throughout the molecular, genetic, and genomic revolutions, but the metagenomic revolution made it impossible to ignore the thousands of understudied species found in every facet of our world (Dewhirst et al. 2010; Quast et al. 2013; Parks et al. 2018). The scientific rise of the microbiome is exciting, but it presents an enormous practical challenge for microbiology. If it took centuries to learn the details of only a few model species, how can we ever understand the thousands of newfound species?

To illustrate the paucity of data on understudied microbes, we performed a bibliometric analysis to quantify the uneven distribution of microbiology research. Release 202 of the GTDB database (Parks et al. 2022) includes 43,409 unique species, and we counted the number of PubMed articles that refer to each species in their title or abstract. The results were heavily skewed. Almost 74% of all known species have never been the subject of a scientific publication—these are *unstudied* bacteria (Figure 1a). Even among the species studied (those with at least one publication), 50% of all articles refer to only ten species (Figure 1b). More than 90% of all bacteriology articles study fewer than 1% of the species, creating a “long tail” of understudied microbes.

**Figure 1:**
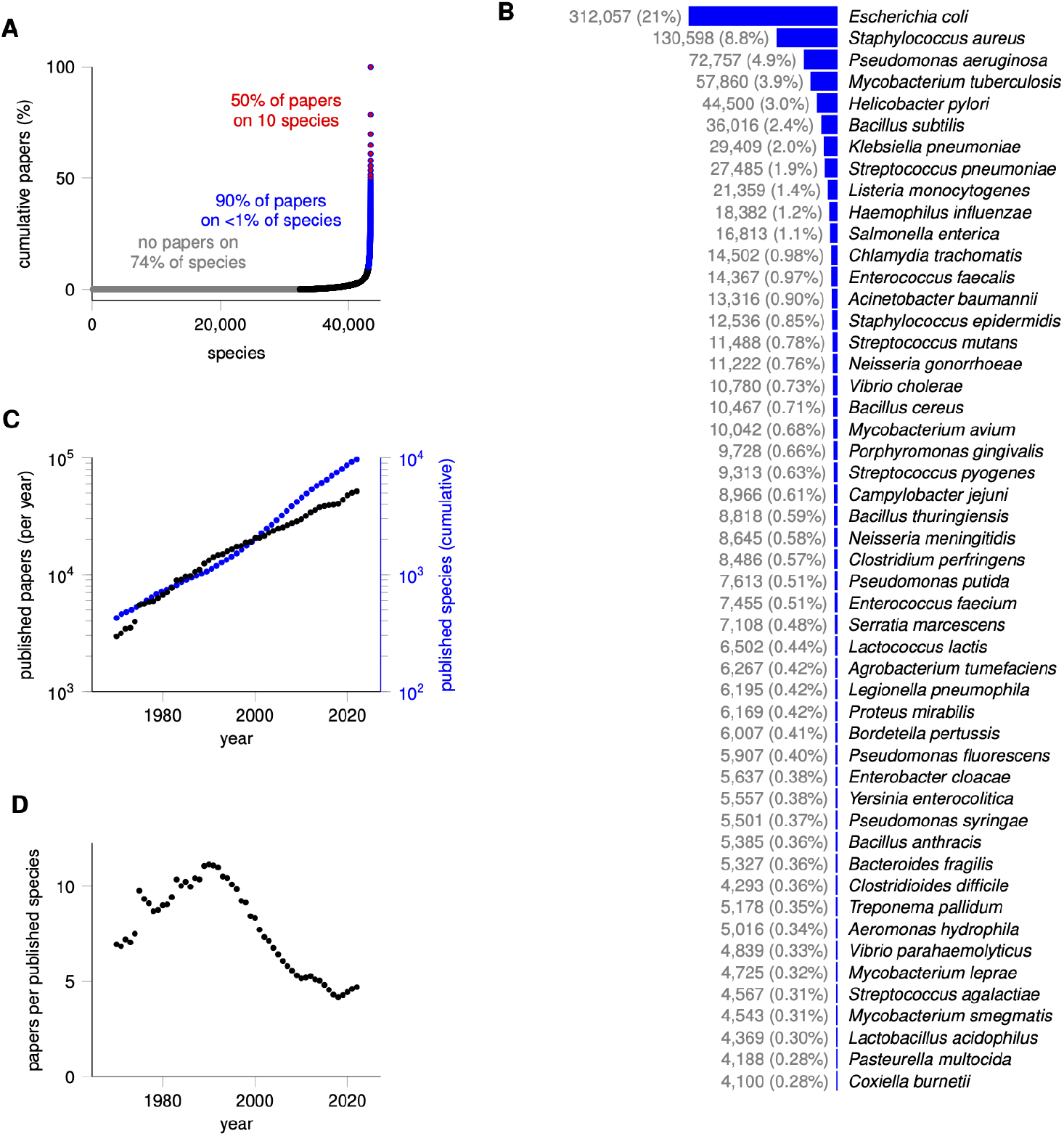
**A.** Most species of bacteria have never been the subject of a scientific paper. We counted how many papers in the PubMed database (up to November 1, 2024) reference on each of 43,409 species of bacteria in their title or abstract. Nearly 74% of species have never been studied (gray dots). The final 1% of most-studied species are the subject of 91.5% of papers, with 10 species appearing in 51.0% of all papers. **B**. Publications are unevenly distributed across the 50 most studied bacteria. The blue bar represents the relative fraction of papers published on each species. The number of articles in PubMed and the percentage of all bacteriology papers published appear to the left. **C**. The number of bacteriology papers published each year (black) grows slower than the cumulative number of species that have been the subject of a paper in PubMed (blue). Note the logarithmic scaling for both papers and species. While the number of papers is increasing, the number of papers *per species* has decreased since 1990, as revealed by panel **D**. It is important to remember that the number of papers published on each species remains highly skewed, and that the plots in **C** and **D** exclude the 74% of species that have never been the subject of a scientific paper.

The scientific enterprise is expanding, and every year scientists publish 4–5% more papers than the previous year (National Science Foundation and National Science Board 2021). It is tempting to think that the increase in scientific output will overcome the long tail of microbes, that is, scientists will eventually get around to studying every species. Unfortunately, the number of species discovered each year outpaces the increases in scientific output (Figure 1c). Between the years 1990–2020, the number of papers published per studied species of bacteria decreased by 60% (Figure 1d). Thus our knowledge density—the amount we learn per species—is actually decreasing.

Our view of bacterial diversity is biased when so much of our understanding comes from so few microbes. Microbiologist Jeffery Gralnick once quipped that “*E. coli* is a great model organism—for *E. coli*.” Gralnick’s comment referenced the discovery of anomalies (relative to *E. coli*) in the TCA cycle of *Shewanella oneidensis* (Brutinel and Gralnick 2012). Although *S. oneidensis* has 201-fold fewer citations that *E. coli*, it is arguably not an understudied species. Our analysis ranks it as the 94th most studied bacterium, which is in the top 2.17% of all species. Even the introduction to Gralnick’s aforementioned paper refers to *S. oneidensis* as a “model environmental organism”. If differences like *S. oneidensis*’ TCA cycle can be found just outside the microbial 2%, imagine the diversity that lies in the other 98% of microbes.

How can microbiologists catch up to the exploding tree of life? We propose two grand challenges for training a generation of microbiologists who can tackle the diversity of the microbial world. First, we need to embrace multifactorial experiment design. There are far too many species, strains, genes, environments, stressors, and phenotypes to study one at a time. Statisticians have taught for decades that the most efficient and robust experimental designs vary multiple factors simultaneously and then deconvolve the effects and interactions with simple statistical models (Fisher 1935). Despite the sound theoretical basis for multifactorial experiments, biologists are routinely taught that “good” experiments vary only a single factor. Teaching multifactorial design would improve the efficiency of microbiologists and illuminate many of the interactions between genes, environments, and cells.

Our second suggestion is to focus on the production of knowledge, not the collection of data. Microbiology is awash in big data, but knowledge production remains bottlenecked by the limited supply of human microbiologists. Statistical and computational tools help distill data for humans, but techniques based on pattern recognition emphasize commonalities between microbes rather than the unique features of each species. The databases themselves present a skewed view of microbial diversity. For example, more than half of the 791 transcriptomics experiments in the BV-BRC database come from five species (Olson et al. 2023). Even if we developed tools to convert all these data into biological knowledge, that knowledge would illuminate a tiny slice of the microbial world. Attempts to repurpose data for new species depend, ironically, on how similar that new species is to our model bacteria. Instead, our knowledge of truly understudied species progresses slowly as scientists publish individual papers using bespoke experiments from their own laboratories. Automating microbiology with robotics and artificial intelligence will accelerate our field (King et al. 2009; Dama et al. 2023), but we need to apply these tools to the myriad species that live in the understudied corners of our world.

Finally, we note that our analysis of microbial diversity includes only a small slice of bacteriology. We did not analyze the literature for viruses, archaea, fungi, or other microbes. Thousands of papers study microbial communities or the microbiome as a whole, but our bibliographic searches only identified papers that named species in their title or abstract. Other papers, such as metagenomic analyses of complex communities, may associate several species with diseases or ecological niches, but our analysis does not capture the species named in the main text, tables, or supplements of these papers. Our study thus reinforces the knowledge gap between microbes that are only studied *en masse* as communities and those select few species whose molecular, genetic, or physiological diversity is studied in detail.

## Methods

Our bibliographic searchers used the NCBI Entrez Direct E-utilities software (Sayers et al. 2024). We searched for each species’ full name (“*Staphylococcus aureus*”) and abbreviated name (“*S. aureus*”). Any subspecies or strain identifiers in the GTDB database were removed before searching, and duplicate species names were removed. Some species share abbreviated names, e.g. *Staphylococcus aureus* and *Streptomyces aureus*; in these cases, we performed separate searches using the full and abbreviated names and kept the only full name results for the species with fewer full name results. Full name searches were also used for species with abbreviated names that form common words, such as *Aminobacterium mobile* = *A. mobile*. These cases were manually identified by their aberrantly high ratio of abbreviated name results to full name results.

All analyses were performed in the R programming language (R Core Team). Visualization were created with pgfplots (Feuersaenger 2020).

## Data and Availability

All code and data used in our analysis is available on our lab website (http://jensenlab.net/publications).

## Funding

This work was supported by the National Institutes of Health (grant GM138210).

## Notes

### Competing Interest Statement

The authors have declared no competing interest.

